# Impact of altering gut microbiota metabolism on osteomyelitis severity in obesity-related type 2 diabetes

**DOI:** 10.1101/2022.01.13.476282

**Authors:** Tina I. Bui, Ann Lindley Gill, Robert A. Mooney, Steven R. Gill

## Abstract

*Staphylococcus aureus* is an opportunistic pathogen causing osteomyelitis through hematogenous seeding or contamination of implants and open wounds following orthopedic surgeries. The severity of *S. aureus*-mediated osteomyelitis is enhanced in obesity-related type 2 diabetes (obesity/T2D) due to chronic inflammation impairing both adaptive and innate immunity. Obesity-induced inflammation is linked to gut dysbiosis, with modification of the gut microbiota by high-fiber diets leading to a reduction in the symptoms and complications of obesity/T2D. However, our understanding of the mechanisms by which modifications of the gut microbiota alter host infection responses is limited. To address this gap, we monitored tibial *S. aureus* infections in obese/T2D mice treated with the inulin-like fructan fiber, oligofructose. Treatment with oligofructose significantly decreased *S. aureus* colonization and lowered proinflammatory signaling post-infection in obese/T2D mice, as observed by decreased circulating inflammatory cytokines (TNF-α) and chemokines (IP-10, KC, MIG, MCP-1, and RANTES), indicating partial reduction in inflammation. Oligofructose markedly shifted diversity in the gut microbiota of obese/T2D mice mice, with notable increases in the anti-inflammatory bacterium, *Bifidobacterium pseudolongum*. Analysis of the cecum and plasma metabolome suggested polyamine production was increased, specifically spermine and spermidine. Oral administration of these polyamines to obese/T2D mice resulted in reduced infection severity similar to oligofructose supplementation, suggesting polyamines can mediate the beneficial effects of fiber on osteomyelitis severity. These results demonstrate the contribution of gut microbiota metabolites to the control of bacterial infections distal to the gut and polyamines as an adjunct therapeutic for osteomyelitis in obesity/T2D.

**Importance:** Individuals with obesity-related type 2 diabetes (obesity/T2D) are at a five times increased risk for invasive *Staphylococcus aureus* osteomyelitis (bone infection) following orthopedic surgeries. With increasing antibiotic resistance and limited discoveries of novel antibiotics, it is imperative we explore other avenues for therapeutics. In this study, we demonstrated that the dietary fiber oligofructose markedly reduced osteomyelitis severity and hyperinflammation following acute implant-associated osteomyelitis in obese/T2D mice. Reduced infection severity is associated with changes in gut microbiota composition and metabolism as indicated by increased production of natural polyamines in the gut and circulating plasma. This work identifies a novel role for the gut microbiome in mediating control of bacterial infections and polyamines as beneficial metabolites involved in improving the obesity/T2D host response to osteomyelitis. Understanding the impact of polyamines on host immunity and mechanisms behind decreasing susceptibility to severe implant-associated osteomyelitis is crucial to improving treatment strategies for this patient population.

## INTRODUCTION

Individuals with obesity-related type 2 diabetes (obesity/T2D) are at increased risk for invasive *S. aureus* infections as a result of obesity-induced chronic inflammation, immune cell dysfunction, and impaired immune defenses (1–4). Osteomyelitis in this patient population is severe, with infections often becoming chronic and requiring secondary surgeries for debridement or replacement of implants, resulting in greater morbidity (5). Obese/T2D individuals undergo total joint arthroplasties at a higher rate than lean, non-diabetic individuals with data from 2011 suggesting more than 80% of patients undergoing joint replacements are obese (6, 7). In general, primary infections following joint replacement surgeries are low (1-3%), but obese/T2D patients have a five-fold increased risk for occurrence of persistent infections in up to 30% of cases (6, 8–10). Furthermore, the majority of joint implant infections are colonized by difficult-to-treat methicillin-resistant *Staphylococcus aureus* (MRSA) (11–14). In an era of decreased antibiotic efficacy due to bacterial resistance, exploring other avenues that target and enhance host response is urgently needed to identify novel, non-invasive therapeutics in this patient population.

Impaired immunity in obese/T2D increases susceptibility to invasive infections. Obese/T2D hosts are immunocompromised, whereby chronic inflammation induced by adipokines such as TNF-α, leads to an accumulation of senescent cells (4, 15). Neutrophils isolated from diabetic rodents and patients are impaired in all the steps of their response: adherence, chemotaxis, phagocytosis, and killing (16–19). Obesity/T2D also modifies T cell function and humoral response to *S. aureus* osteomyelitis (20, 21). Elucidation of functions in obesity/T2D that contribute to inflammation and exacerbate *S. aureus* osteomyelitis will improve understanding of the effects of inflammation on infection outcomes (4).

Dysbiosis of the gut microbiome in obesity/T2D is a significant contributor to chronic inflammation and insulin resistance (22, 23). Gut dysbiosis is influenced to a greater extent by obesity than diabetes in obese/T2D patients, with significant changes in host metabolism (24). In obesity-related dysbiosis, overall diversity of the gut microbial community is reduced along with compositional changes leading to higher energy harvest from diet and subsequent total body fat accumulation (22). Compelling evidence suggests that disruption of the gut microbial community by a high-fat diet leads to increased inflammation in obesity/T2D (25–27). This shift in the gut microbiota of obese/T2D rodents and individuals is linked to increased circulating lipopolysaccharide (LPS), a bacterial cell wall component recognized by toll-like receptor 4 (TLR4) (26–28). Persistent activation of TLR4 presented on innate immune cells triggers release of proinflammatory cytokines which promote chronic inflammation and other complications of obesity.

The obesity-induced gut dysbiosis is reversed by changes in the host diet, a primary driver of gut microbiota composition and metabolism (25). One way to modulate the obese/T2D gut microbiota is through indigestible dietary fiber composed of fermentable carbohydrates (29). Mammalian hosts do not possess enzymes to hydrolyze these carbohydrates, and can only process and absorb nutrients after they are catabolized by gut microbes. Prior studies exploring the effects of an inulin-like fructan, oligofructose, have demonstrated its importance in improving systemic complications of obesity/T2D including weight loss, glycemic control, insulin resistance, and inflammation (30–34). Fermentation of oligofructose by gut microbes in the large colon produces numerous metabolic products, including short-chain fatty acids (SCFAs), which affect a broad range of cellular and immune functions (35–37). In our study, obese/T2D mice with tibial *S. aureus* infections are treated with oligofructose to reverse gut dysbiosis and reduce inflammation which exacerbates infections. We hypothesized that alteration of the obese/T2D gut microbiota would lead to improvements in osteomyelitis infection outcomes through changes in bacterial metabolite production. Our results demonstrate that oligofructose reduced bone infection severity in obese/T2D mice, through alterations in the gut microbiota and downregulating inflammation 14-days post-infection. Importantly, we identified a novel role for polyamines as beneficial metabolites that directly improved infection outcomes similar to oligofructose.

## RESULTS

### Oligofructose decreases *S. aureus* implant-associated infection severity in obese/T2D mice

Oligofructose (OF) is a soluble fiber which has been reported to alleviate some of the secondary complications of obesity and type 2 diabetes in humans and rodents (30–34). To determine if oligofructose has an effect on implant-associated osteomyelitis severity in obese/T2D mice, we utilized a diet-induced murine model of obesity/T2D and supplemented lean and high-fat diets with oligofructose for two weeks prior to and during infection. **(Fig 1A)**. The insoluble fiber cellulose (CL) was used as a control supplement that is poorly fermented by gut microbes and resistant to mammalian metabolism. Results from a glucose tolerance test performed prior to tibial transplant at week 14 demonstrated that obese/T2D+CL control mice exhibit increased glucose tolerance with a modest improvement as seen in obese/T2D+OF mice **(Fig S1)**. After initiation of a tibial implant-associated *S. aureus* USA300 infection, infections were monitored for 14 days, a time-frame within which acute osteomyelitis can develop. In agreement with previous research suggesting obese/T2D mice are at increased risk for more severe invasive osteomyelitis compared to healthy counterparts, the obese/T2D+CL control mice exhibited more severe infections than lean/control mice regardless of supplement, with 5% greater weight loss **(Fig 1B)** and increased abscess formation and bacterial burden **(Fig 1C-E)** (38, 39). Oligofructose reduced soft-tissue abscess size by ~50% **(Fig 1C and 1D)** and the associated bacterial burden in the abscess and tibia following *S. aureus* implant-associated infections by ~1 log fold as compared to the obese/T2D+CL control mice **(Fig 1E)**. The reduction in both measures of infection with oligofructose was only observed in obese/T2D mice and not lean/control mice. These results demonstrate that oligofructose was specifically effective in reducing infection severity within the obese/T2D subject group and therefore has therapeutic potential.

**Figure 1.**
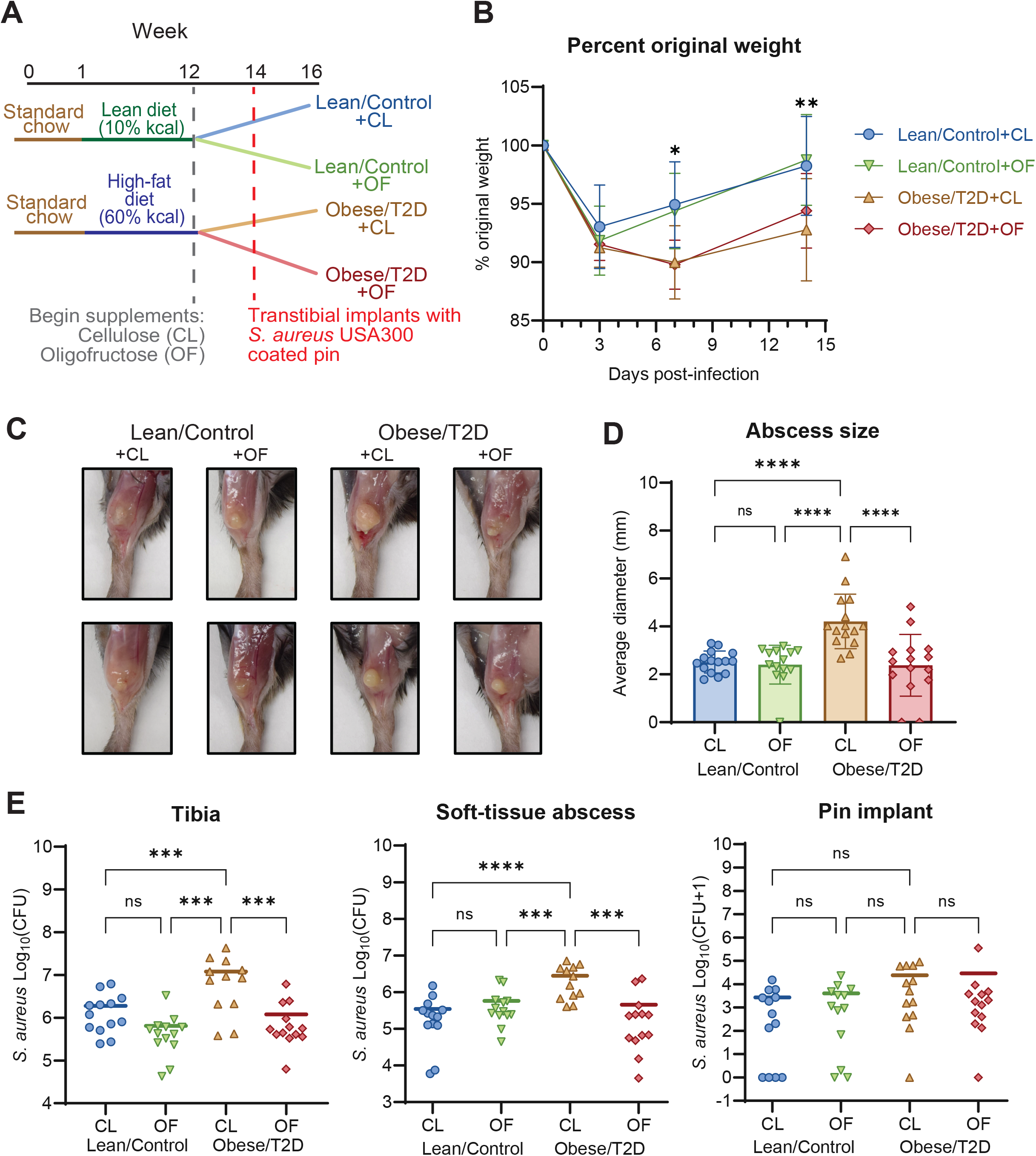
Oligofructose decreases *S. aureus* burden during implant-associated osteomyelitis in obese/T2D mice. A) Schematic of the 16-week experimental timeline. CL=Cellulose (control fiber); OF=Oligofructose. Lean/control and obese/T2D refers to the disease state and CL (cellulose, control fiber) or OF (oligofructose, experimental fiber) refers to the dietary supplement. For example, lean/control treated with cellulose is denoted as lean/control+CL while control obese/T2D treated with cellulose is obese/T2D+CL. B) Percent original weight of mice during the course of infection. Significance between lean/control+CL and obese/T2D+CL (* *P* < 0.05, ** *P* < 0.01) by two-way ANOVA with Tukey’s post-hoc test. There were no differences between obese/T2D+CL and obese/T2D+OF. C) Two representative images of soft-tissue abscesses from each treatment group 14 days post-infection. D) Quantification of abscess size using digital caliper. E) Infected soft tissue, tibias, and pins were collected at day 14 post-infection for enumeration of *S. aureus* with bars representing means. Bar graphs represent mean ± SD. n=13. Significance was identified using one-way ANOVA and Tukey’s post-hoc multiple comparisons test. *\*\*P*<0.01, *\*\*\*P*<0.001, *\*\*\*\*P*<0.001.

### Oligofructose reduces *S. aureus* community colonization in the bone

Spatial colonization of *S. aureus* in the bone marrow and bone osteolysis was examined as an additional outcome of osteomyelitis severity. Infected tibias were sectioned and stained with 1) Brown-Brenn Gram stain to visualize *S. aureus* communities in crystal violet, 2) alcian blue hematoxylin-orange G (ABHO) to visualize mature calcified bone orange to red and soft tissues pink to red for abscesses, and 3) tartrate-resistant alkaline phosphatase (TRAP) for osteoclast activation in pink to indicate areas of osteolysis **(Fig 2A)**. Histological analysis showed that while the number of abscesses were similar across treatment groups **(Fig 2B)**, the total areas colonized by *S. aureus* in an abscess community or in a biofilm associated with necrotic bone differed. Consistent with bacterial CFU quantification of the bone **(Fig 1E)**, we observed ~60% increased colonization of *S. aureus* communities in the bone marrow of obese/T2D+CL control mice as compared to obese/T2D+OF and lean/control mice when normalizing positive Gram stain areas to whole tibia area **(Fig 2C)**. Analysis of osteoclast activation by TRAP staining showed no differences in osteolysis across groups **(Fig 2D)**. This corresponded with µ-CT imaging of the implant site **(Fig 2E)** and quantification of the infected hole size for osteolysis **(Fig 2F and 2G)**. Together, these data suggested that obesity/T2D may not impact osteolysis during acute osteomyelitis (14 days post-infection). Importantly, oligofructose modestly reduced *S. aureus* community colonization in the infected bone.

**Figure 2.**
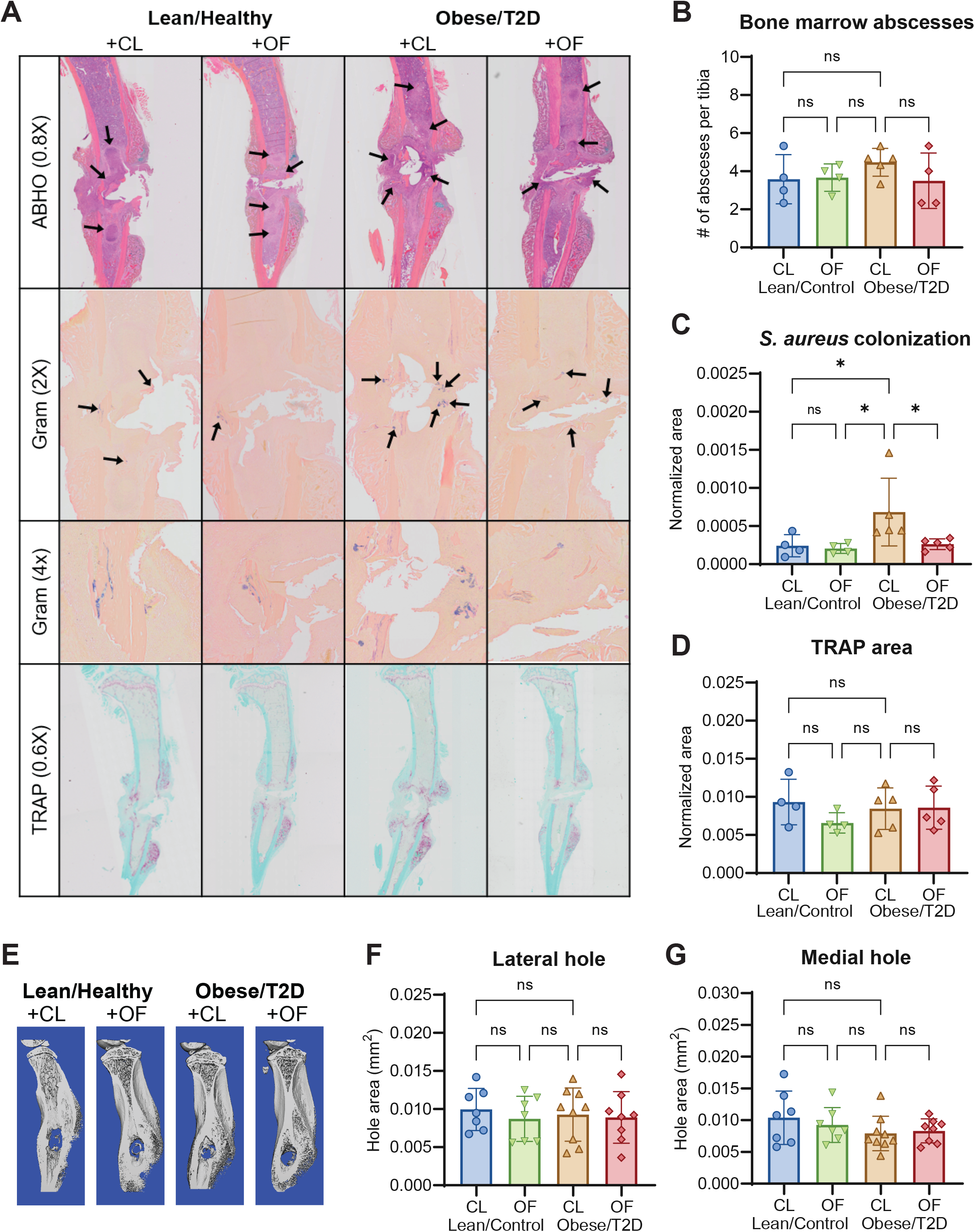
Oligofructose decreases *S. aureus* colonization in the bone marrow. A) Histologic sections of infected tibias at 14 days post-infection were stained with alcian blue hematoxylin orange G (ABHO) for visualization of abscess formation, Brown-Brenn Gram-stains for *S. aureus* colonization, and tartrate-resistant acid phosphatase (TRAP) for osteoclast detection. B) Quantification of bone marrow abscesses by counting across three sections per mouse. C) Total *S. aureus* community colonization area in the bone marrow was quantified by using an analysis protocol package (APP) in Visiopharm which identified positive Gram-stained area and normalized to total tibial area. At least three sections were measured per mouse. D) TRAP area was also quantified by an APP in Visiopharm and normalized to total tibial area. At least three sections were measured per mouse E) Representative images of μCT analyses on infected tibias harvested 14 days post-infection. F and G) Bar graphs represent mean ± SD. Histology n=7-8, µCT n =5. Significance was identified using one-way ANOVA and Tukey’s post-hoc multiple comparisons test. (\**P < 0*.*05*). CL=Cellulose (control fiber); OF=oligofructose.

### Oligofructose downregulates inflammatory signals in obese/T2D mice following infection

It is well established that the host mounts a temporary inflammatory response during acute osteomyelitis. However, chronic inflammation combined with infection in obesity/T2D can lead to hyperinflammatory responses manifesting in the form of increased immune cell recruitment and cytokine storm which then exacerbate infection outcomes (40–43). To examine the effects of oligofructose on immune signaling associated with reduced infection severity, we next determined the levels of cytokines at 14 days post-infection. The obese/T2D+CL control mice displayed a hyperinflammatory immune response with overall increases in cytokines and chemokines post-infection that were not present in other treatment groups **(Fig 3A)**. We observed a sustained production of TNF-*α* and IL-6 cytokines at 14 days post-infection in the obese/T2D+CL control that was absent in both lean/control groups and obese/T2D+OF mice **(Fig 3B and 3C)**. Consistent with this reduction in inflammation, obese/T2D+OF mice showed a markedly decreased expression of several chemokines as compared to obese/T2D control mice, including KC (CXCL1), MCP-1 (CCL2), MIG (CXCL9), IP-10 (CXCL10), and RANTES (CCL5) **(Fig 3D-H)**. MIG and IP-10 are recruitment signals produced when induced by IFN-γ, both targeting activated T cells via chemokine receptor 3 (44). KC and MCP-1 primarily target innate immune cells, including neutrophils and monocytes respectively (45–47). From these results, we conclude that oligofructose reversed hyperinflammation that is associated with severe osteomyelitis and higher morbidity in control obese/T2D mice.

**Figure 3.**
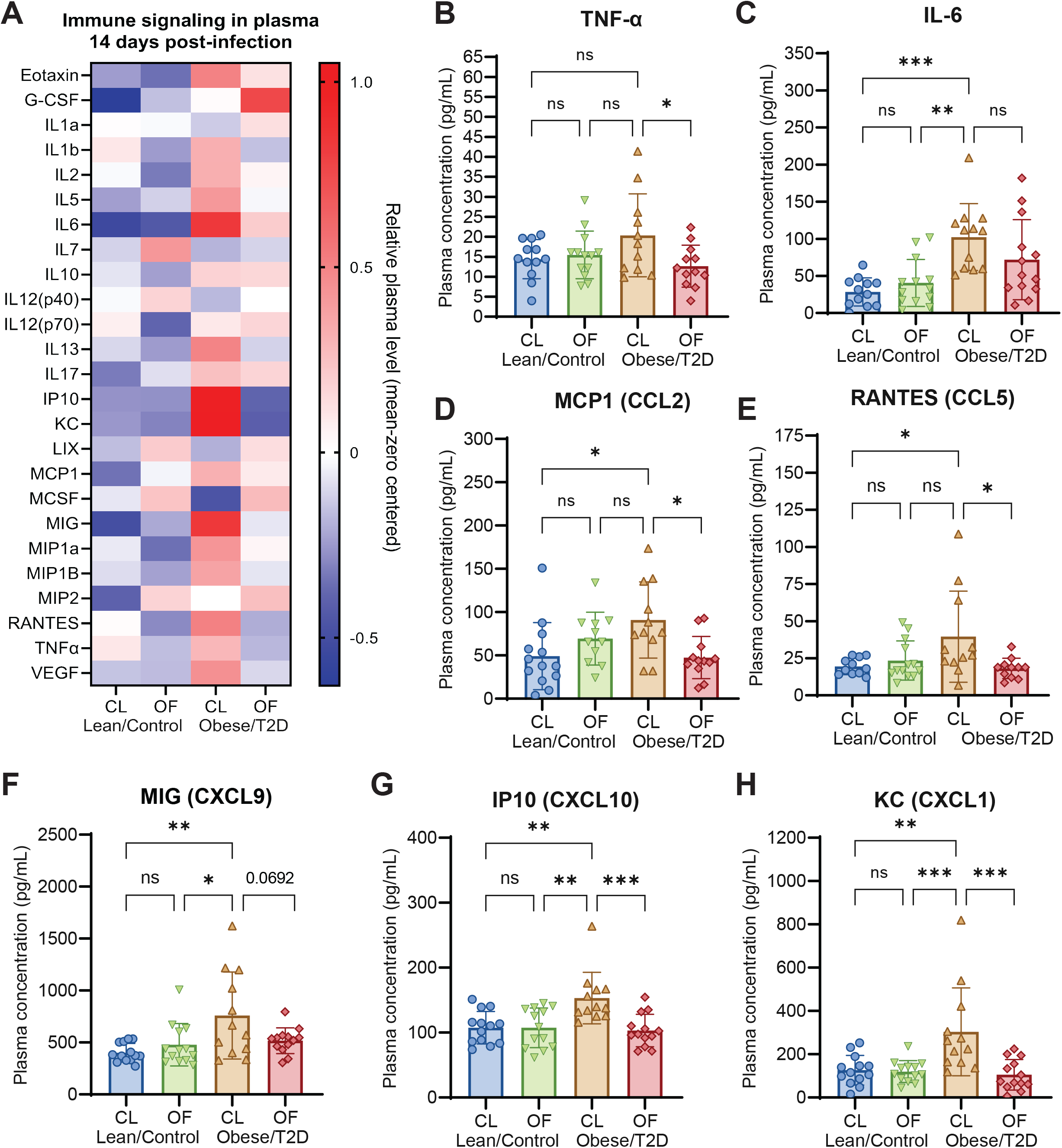
Oligofructose downregulates inflammatory signals in obese/T2D mice following infection. A) Heat-map of mean-zero centered concentrations of different cytokines/chemokines analyzed from plasma isolated 14 days post-infection. B-H) Significantly altered cytokines and chemokines. Bar graphs represent mean ± SD. n=13. Significance was identified using one-way ANOVA and Tukey’s post-hoc multiple comparisons test. (*\*P* <0.05, *\*\*P* < 0.01, *\*\*\*P* < 0.001). CL=Cellulose (control fiber); OF=oligofructose.

### Oligofructose shifts obese/T2D gut microbiota away from obesity-related dysbiosis

We performed 16S rRNA amplicon sequencing of mouse fecal samples collected weekly, from weeks 0 through 16, to identify taxonomic shifts in community composition associated with reduced infection severity in response to oligofructose. All mice on standard chow at week 0 had similar microbiota composition prior to the dietary regimen **(Fig 4A)**. In agreement with prior studies, obese/T2D mice on the high-fat diet with no supplement (obese/T2D+NS) exhibited gut dysbiosis, with a shift of the microbiota to a taxonomic composition distinct from lean/control+NS mice throughout weeks 1-12. Lean/control+CL and obese/T2D+CL mice during weeks 13-16 had gut microbiota composition similar to that observed prior to supplementation at week 12 **(Fig 4B)**. However, following supplementation with oligofructose through weeks 13-16, the obese/T2D+OF microbiota was distinct from all other groups **(Fig 4B)**. These results collectively demonstrate that oligofructose markedly altered the obese/T2D gut microbiota, while cellulose had little effect and served as a negative control for fiber supplementation.

**Figure 4.**
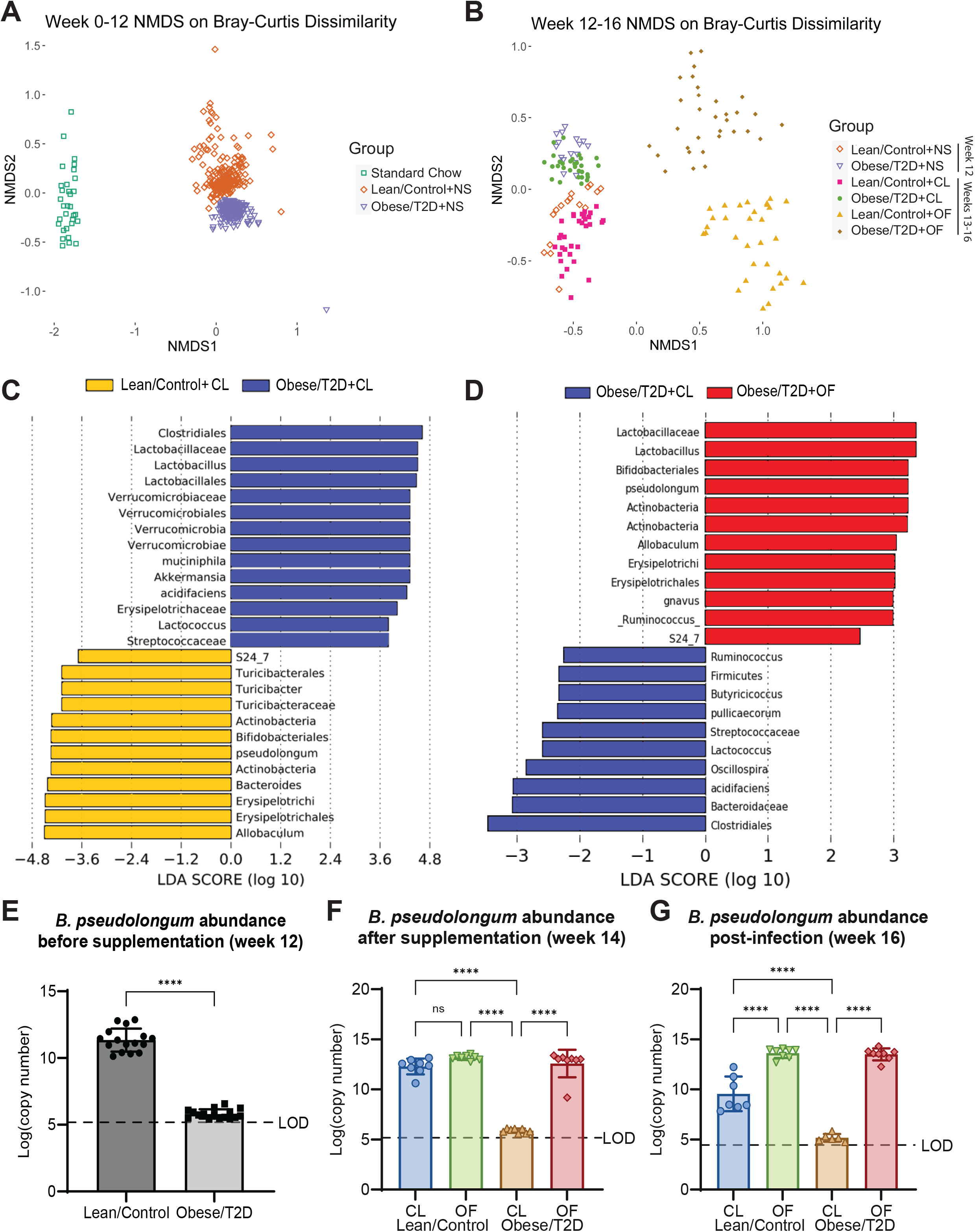
Oligofructose shifts gut microbiota away from diseased state by inducing the expansion of *Bifidobacteria pseudolongum*. A) Nonmetric multidimensional scaling (NMDS) plot of Bray-Curtis dissimilarity metric during week 0-12. NS=no supplement B) NMDS plot of Bray-Curtis dissimilarity during week 12-16. Week 12 represents mice prior to supplementation (NS=no supplement). Week 13-14 represent mice supplemented with CL=Cellulose (control fiber) or OF=oligofructose, before infection and during infection week 15-16. C and D) Pairwise analysis using LEfSe was used to identify OTUs discriminating between two indicated groups. LDA scores were calculated based on a subset of vectors consisting of Kruskal-Wallis tests analyzing all features and pairwise Wilcoxon test checks between groups. E-G) Absolute abundance of *B. pseudolongum* identified by qPCR for weeks 12, 14, and 16. Bar graphs represent mean ± SD. LOD = Limit of detection. n=7-8. Significance was identified using one-way ANOVA and Tukey’s post-hoc multiple comparisons test. (\*\*\*\**P*<0.0001).

To identify operational taxonomic units (OTUs) discriminative of each group, we performed Linear discriminant analysis Effect Size (LEfSe) on the gut microbiota at 14 days post-infection (week 16). Several taxa were differentially abundant between lean/control+CL and obese/T2D+CL control mice **(Fig 4C)**. The gut microbiota of obese/T2D+CL control mice was more closely associated with *Akkermansia muciniphila* and *Lactobacillus* spp **(Fig 4C)**. Multiple *Lactobacillus* species have been associated with weight gain and a fiber-deprived diet (48–50). In contrast, lean/control+CL mice were associated with *Bifidobacterium pseudolongum*, an anti-inflammatory bacterium that produces immunodulatory short-chain fatty acids (SCFAs) from fiber fermentation and an unknown species of S24-7 family and *Allobaculum* genus **(Fig 4C)** (51, 52). Oligofructose treatment in obese/T2D+OF mice also led to increased abundance of *B. pseudolongum* and unknown taxa from the S24-7 family and *Allobaculum* genus **(Fig 4D and S2)**. Prior to supplementation at week 12, *Bifidobacteria* was present in lean/control mice at an average of 8% relative abundance compared to 0.01% in obese/T2D mice **(Fig S2A)**. Oligofructose supplementation at week 14 rapidly increased *Bifidobacteria* abundance to 34% in lean/control+OF and 18% in obese/T2D+OF mice **(Fig S2B)**. This expansion in *Bifidobacteria* continued while mice were given oligofructose 14 days after infection at week 16, where *Bifidobacteria* abundance was found to be at 44% and 39% lean/control+OF and obese/T2D+OF mice respectively **(Fig S2C)**. We next quantified absolute abundance of *B. pseudolongum* for weeks 12, 14 and 16 by qPCR **(Fig 4E-G)**. Without any dietary fiber supplement at week 12, obese/T2D mice had 6-log fold less *B. pseudolongum* than lean/control mice with values near the detection limit **(Fig 4E)**. After oligofructose supplementation but prior to infection (week 14), the copy number of *B. pseudolongum* in obese/T2D+OF increased to levels similar to that of lean/control mice, **(Fig 4F)**. Following 14 days of infection, *B. pseudolongum* abundance remained similar to week 14. Overall, these results illustrate that oligofructose induces large, distinct shifts in the gut microbiota of obese/T2D mice away from obesity-induced dysbiosis and more similar to that of lean/control mice. The considerable changes in the abundance of *B. pseudolongum* and potential contribution to immunomodulation suggested a key role for this bacterium in the microbiota response of obese/T2D mice to dietary oligofructose and a reduction in infection severity associated with increased levels of SCFAs.

### Polyamine production is increased in response to dietary oligofructose

To determine which gut-microbial derived metabolite(s) were altered following oligofructose supplementation, we performed targeted metabolomics on plasma and cecal material isolated from mice after two weeks on supplement but before infection **(Fig 1A)**. Samples were harvested prior to surgery in order to identify potential biomarkers of improved infection outcomes and to prevent active infections from confounding metabolite production. In the plasma, lean/control+CL and obese/T2D+CL control mice had many differentially altered metabolites including the expected elevation of glucose in obese/T2D mice **(Fig 5A)**, which is in agreement with our glucose tolerance test **(Fig S1)**. We unexpectedly observed no differences in abundance of gut and plasma SCFAs **(Fig S3)**. When comparing plasma of obese/T2D+OF to obese/T2D+CL mice, a small decrease in glucose and an approximately 7-fold change in acetyl-ornithine was observed **(Fig 5B)**. Acetyl-ornithine, an intermediate involved in the final production of polyamines through synthesis of arginine and ornithine **(Fig 5C)**, was found to be significantly upregulated in both lean/control+OF and obese/T2D+OF **(Fig 5D)**. Polyamines, which are produced by both the host and gut microbes from arginine and ornithine precursors **(Fig 5C)**, are integral to host cellular processes including apoptosis, proliferation, and differentiation (53). Of the three most common polyamines putrescine, spermidine, and spermine, only spermine and spermidine were found to be elevated 3-4 times in obese/T2D mice fed oligofructose **(Fig 5E)**. Because acetyl-ornithine is microbiota-derived and is used during anabolism of polyamines, increases in acetyl-ornithine following oligofructose supplementation in both lean/control and obese/T2D mice suggest the increases in spermine and spermidine are likely derived from the microbiota.

**Figure 5.**
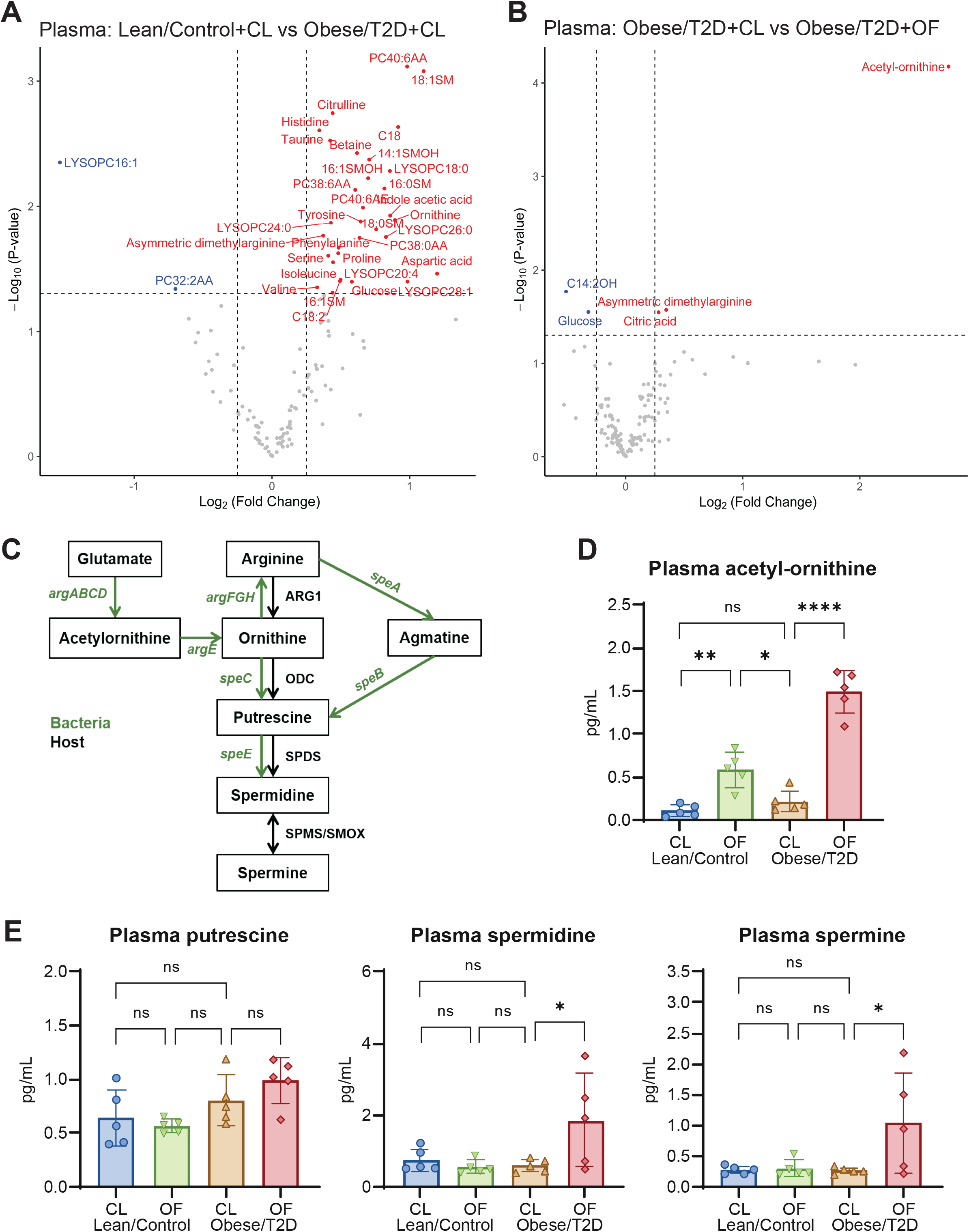
Targeted metabolomics of plasma. A and B) Volcano plots of metabolomic profiles of plasma were compared between Lean/Control+CL vs Obese/T2D+CL and Obese/T2D+CL vs Obese/T2D+OF. Metabolites annotated in blue and red are significantly decreased and increased respectively. C) Mammalian and bacterial biosynthetic pathway for polyamines. ARG1 (arginase 1), ODC (ornithine decarboxylase), SPDS (spermidine synthase), SPMS (spermine synthase), SMOX (spermine oxidase). D) Absolute concentration of the polyamine precursor acetyl-ornithine. E) Absolute concentrations of the three natural polyamines detected. Bar graphs represent mean ± SD. n=5. Significance was identified using one-way ANOVA and Tukey’s post-hoc multiple comparisons test. (*\*P* <0.05, *\*\*P* < 0.01, *\*\*\*\*P* < 0.0001). CL=Cellulose (control fiber); OF=oligofructose.

Similar trends were observed in analysis of metabolites from the cecum **(Fig 6A and 6B)**. Both spermine and acetyl-ornithine were upregulated in obese/T2D+OF compared to obese/T2D+CL controls **(Fig 6B)**. When examining absolute concentrations of metabolites in the polyamine biosynthesis pathway, acetyl-ornithine increased by 8-fold and spermine approximately quadrupled in obese/T2D+OF mice as compared to obese/T2D+CL control mice, while spermidine levels doubled but were not statistically significant **(Fig 6C)**. These results suggest that altered biosynthesis of polyamines as a result of oligofructose supplementation and shift in the microbiota, may contribute to a reduction in infection severity.

**Figure 6.**
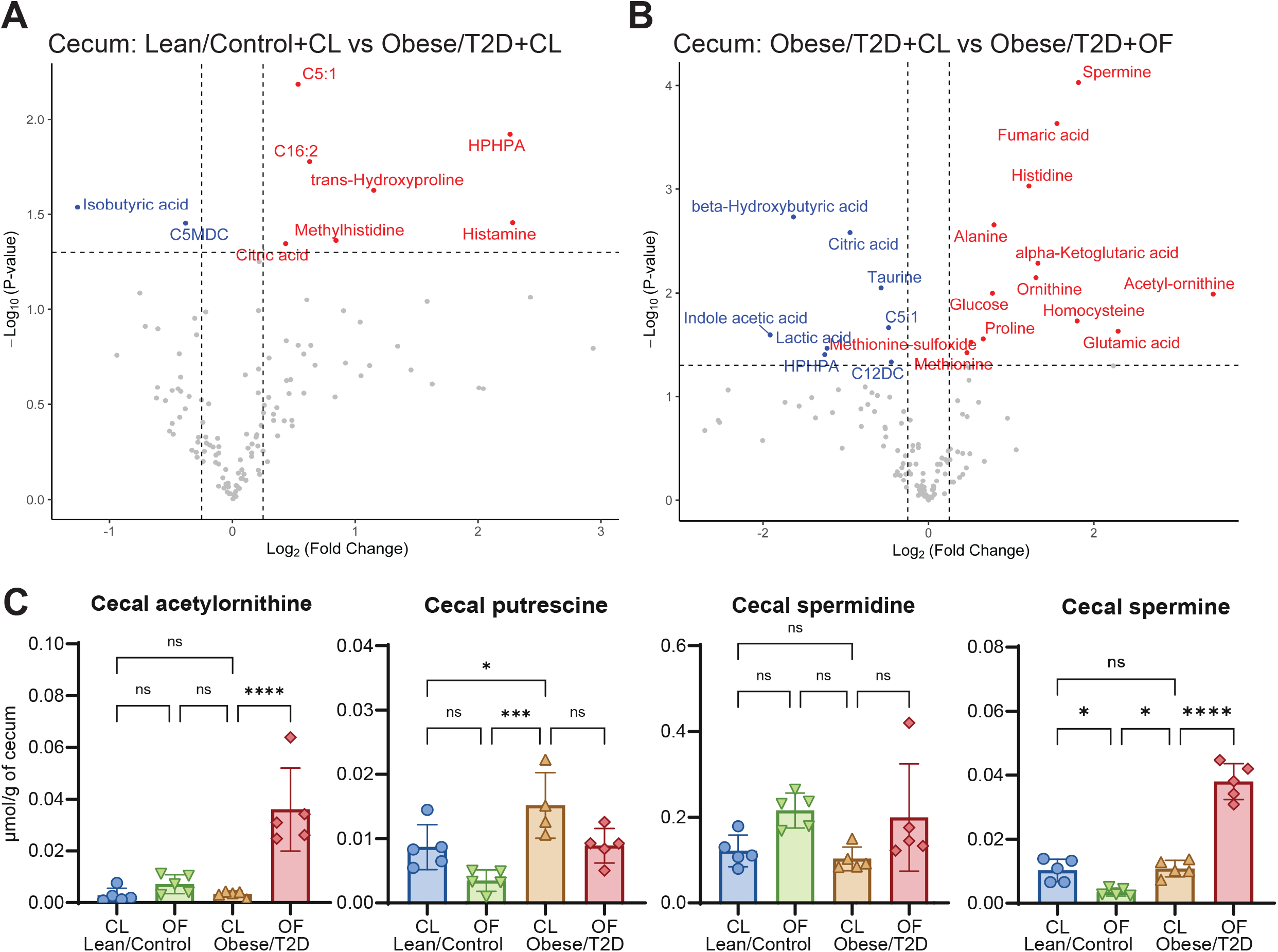
Targeted metabolomics of cecum. A and B) Volcano plots of metabolomic profiles of cecal material were compared between Lean/Control+CL and Obese/T2D+CL and Obese/T2D+CL vs Obese/T2D+OF. Metabolite concentrations were normalized based on grams of cecal material. Metabolites annotated in blue and red are significantly decreased and increased respectively. C) Normalized concentrations of targeted metabolites involved in polyamine biosynthesis. Bar graphs represent mean ± SD. n=5. Significance was identified using one-way ANOVA and Tukey’s post-hoc multiple comparisons test. (*\*P* <0.05, *\*\*\*P*<0.001). CL=Cellulose (control fiber); OF=oligofructose.

### Oral administration of polyamines reduces infection severity similar to oligofructose in obese/T2D mice

In order to directly determine the effects of polyamines on osteomyelitis severity, mice were supplemented with spermine and spermidine in drinking water two weeks prior to and continuing after infection. Water consumption in mice given polyamines (PA) and controls given PBS was similar **(Fig 7A)**. Weight loss in obese/T2D mice was significantly more than lean/control mice at 3 and 7 days post-infection regardless of chemical supplement **(Fig 7B)**. At day 14, obese/T2D+PA mice had markedly reduced weight loss by 5% compared to obese/T2D+PBS control mice, suggesting efficacy of polyamines in overall recovery. Polyamine treatment in obese/T2D+PA mice also led to 50% smaller abscesses but had no effect in lean/control+PA mice. More importantly, polyamine supplementation decreased the elevated *S. aureus* bone and soft tissue burden in obese mice by approximately 1-log fold while having no effect in lean/control mice. These results demonstrate that the effects of polyamines on *S. aureus-*osteomyelitis in obese/T2D mice were similar to that of oligofructose **(Fig 1)**. Furthermore, the *S. aureus* USA300 LAC::lux strain that was used in our study carries, *speG*, a gene in the arginine catabolic mobile element which confers resistance to polyamines (54, 55). Our *in vitro* growth experiments confirmed that *S. aureus* USA300 LAC::lux is not susceptible to polyamines **(Fig S4)**. Therefore, we conclude that polyamines contribute to the reduced infection severity observed in obese/T2D mice when supplemented with oligofructose through immune modulation **(Fig 8)**.

**Figure 7.**
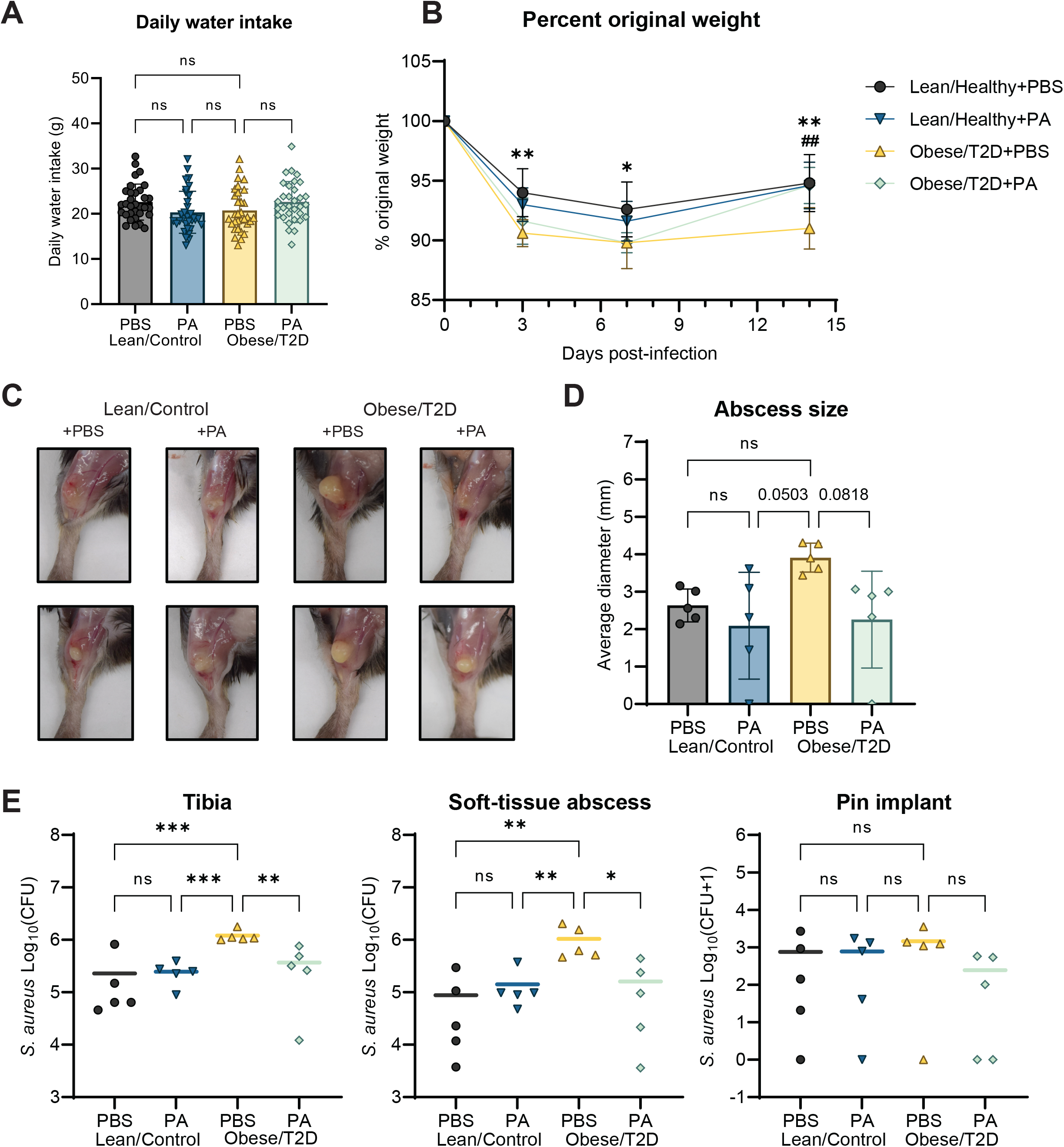
Polyamines reduce infection severity similar to the fiber oligofructose. A) Daily water intake was determined by measuring changes in weight of drinking bottles. B) Percent original weight of mice over 14 days of infection. Significance between lean/control+CL and obese/T2D+CL (* *P* < 0.05, ** *P* < 0.01) and obese/T2D+CL vs obese/T2D+OF (## P < 0.01) was determined by two-way ANOVA with Tukey’s post-hoc test. C) Two representative images of soft-tissue abscesses 14 days post-infection. D) Quantification of size of soft-tissue abscess size seen in C. E) Infected soft-tissue abscesses and bones were homogenized and serially diluted for bacterial quantification 14 days post-infection, with bars representing means. Bar graphs represent mean ± SD. n=5. Significance was identified using one-way ANOVA and Tukey’s post-hoc multiple comparisons test. (*\*P* <0.05, *\*\*P* < 0.01, *\*\*\*P* < 0.001). PBS=phosphate buffered saline (control); PA=polyamines (3mM total spermine+spermidine).

**Figure 8.**
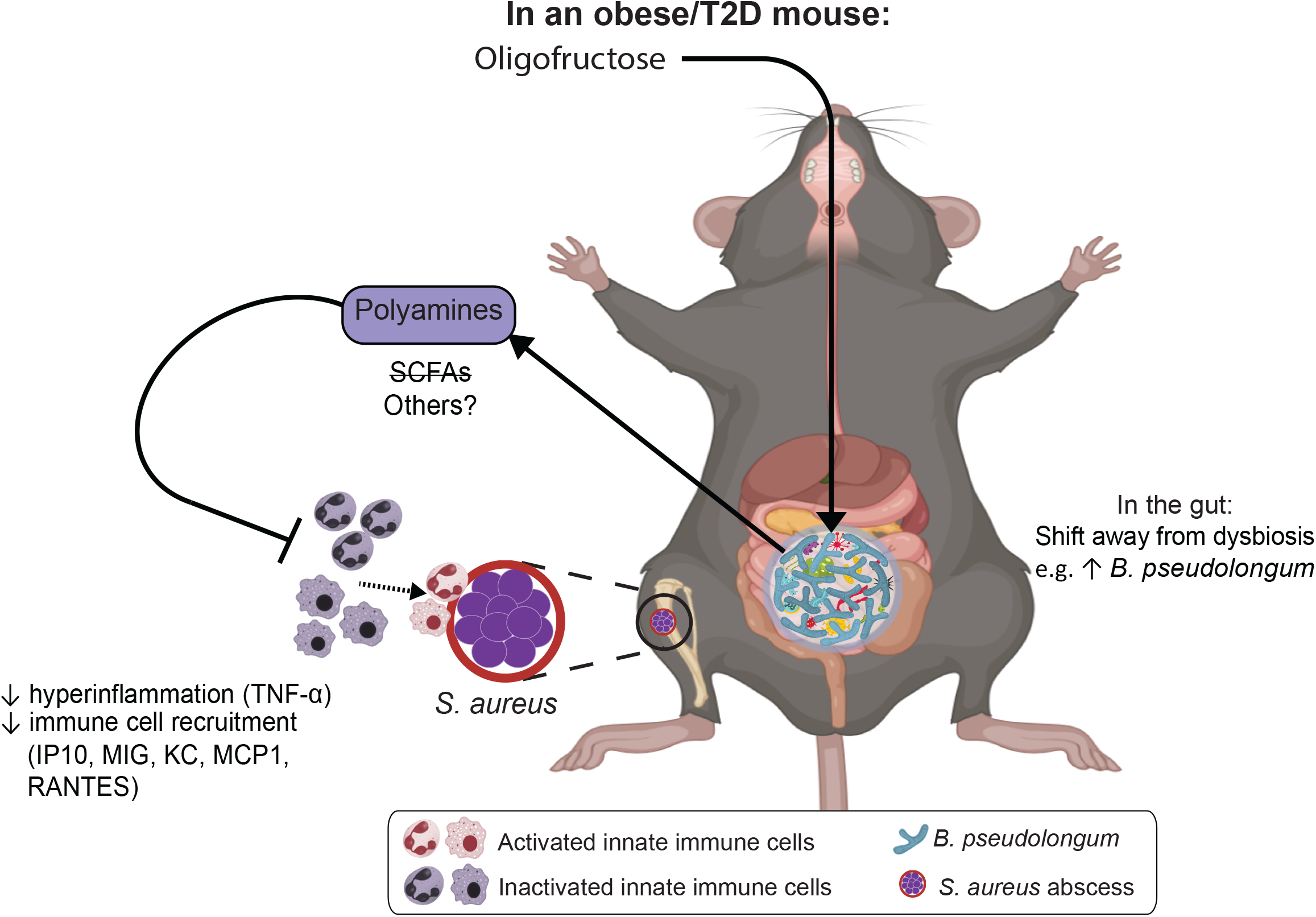
Proposed mechanism mediating the beneficial effects of oligofructose on osteomyelitis in obese/T2D mice. In an obese/T2D mouse, oligofructose supplementation led to increased *Bifidobacteria pseudolongum* in the gut and significantly shifted the gut microbiome away from a non-obese/diabetic state. This was associated with elevated levels of polyamines in the local gut environment and systemically in plasma. Oligofructose supplementation in obese/T2D also resulted in reduced infection severity and reduced pro-inflammatory signaling, suggesting a dampening of hyperinflammatory response seen in control obese/T2D mice. Direct administration of polyamines to obese/T2D mice led to similar improved infection outcomes to oligofructose, suggesting polyamines are involved in modulating the obese/T2D immune responses against *S. aureus-*mediated osteomyelitis.

## DISCUSSION

A major component of host response to infections and systemic inflammation is the gut microbiota (56, 57). In obesity/T2D, gut dysbiosis contributes to chronic inflammation which causes deficits in the immune system, thereby increasing susceptibility to and exacerbating infections. Thus, modulating the gut microbiota of obesity/T2D can potentially identify microbial pathways important for enhancing host immune responses against *S. aureus-*mediated osteomyelitis. One potential modifier of gut dysbiosis is dietary fiber. To explore this potential approach, we supplemented diet-induced obese/T2D mice with an inulin-derivative, oligofructose, and examined infection outcomes including bacterial burden, immune signaling, and metabolic profiles. To our knowledge, this is the first study to show efficacy of oligofructose in reducing infection severity during tibial mplant-associated osteomyelitis, a site distal to the gut. Our results further showed that polyamines were involved in mediating improved infection outcomes.

We demonstrated that oligofructose decreased bacterial burden in both infected tibia and soft tissue of obese/T2D mice as compared to controls **(Fig 1E)**. In contrast, oligofructose did not impact soft-tissue abscess size or bacterial burden in lean/control+OF mice as compared to those in lean/control+CL given cellulose, the fiber control **(Fig 1D and 1E)**. Since lean/control mice without any supplement did not have gut dysbiosis nor the associated inflammation of obesity/T2D **(Fig 3A)**, the beneficial effects of oligofructose may not alter the course of infection as lean/control mice already exhibit normal inflammatory responses. Bacterial burden on the pin implants was also not affected by oligofructose supplementation, likely due to implants acting as fomites harboring *S. aureus* in a biofilm state which physiologically different than abscesses in tissue and bone (58). Oligofructose also decreased the elevated total *S. aureus* community colonization in obese/T2D mice **(Fig 2C)**. Although prior studies have shown obese/T2D mice to have a significant impact on osteolysis, we did not observe an effect in this study **(Fig 2C and 2E-G)**, likely due to measurement at 14 vs 21 days (38, 39). Our model examined the effects of oligofructose during acute osteomyelitis, an infection that causes short-term inflammation, whereas 21 days post-infection emphasizes chronic infections with distinct characteristics such as a fibrotic marrow and periosteal reactive bone formation (59). Nonetheless, we conclude that oligofructose had beneficial effects on infection outcomes in obese/T2D mice and warrants further investigation as a positive modifier of immunity and gut dysbiosis.

In this study, oligofructose prevented sustained inflammation in obese/T2D mice. Several pro-inflammatory signals that were elevated in the obese/T2D control mice were down-regulated by oligofructose post-infection including TNF-α and IL-6 and several chemotactic signals **(Fig 3)**. During acute infections as this model illustrates, inflammation is normally reduced so that wound healing can occur. Therefore, decreased recruitment signals for inflammatory mediators by oligofructose likely reduced tissue damage induced by inflammation, allowing for better wound healing in obese/T2D murine hosts. Additional studies to support this premise include quantification of tissue damage markers such as lactate dehydrogenase or M2 polarization of macrophages, which are associated with inflammation resolution. Nonetheless, the elevated inflammatory signals in obese/T2D+CL control mice suggested a hyperinflammatory response to infection that may contribute to increased infection severity in this patient population. Several other studies have suggested that a hyperinflammatory response can worsen infection outcomes. Nielsen and colleagues demonstrated that obesity-related diabetes exacerbated *Acetinobacter baumanii*-induced sepsis, which improved following administration of the immunosuppressing agent dexamethasone (40). Genetically altered obese/T2D (*db/db*) mice demonstrated increased neutrophil infiltration in infections of the hind paw and heightened production of chemokines CXCL1 and CXCL2 12 hours after infection, suggesting a hyperinflammatory response (41). However, *ex vivo* experiments with peripheral neutrophils isolated from *db/db* mice showed diminished bacterial killing when challenged with *S. aureus* (41). Previous work from our group has also led to similar conclusions. We have reported that diet-induced obese/T2D mice exhibit a hyperinflammatory response with significantly more macrophages recruited to the site of infection corresponding with increased bone destruction and bacteria survival (38). This current study not only provides additional evidence that hyperinflammatory responses seen in obese/T2D hosts are linked to more severe infections, but also evidence for oligofructose moderating inflammation that would normally exacerbate acute osteomyelitis.

To determine a potential mechanism responsible for reduced inflammation and infection severity, we investigated longitudinal changes in the gut microbiota and metabolite profiles in all four groups **(Fig 1A)** of mice. The gut microbiota of obese/T2D mice without any supplement, shifted to a dysbiotic composition distinct from lean/control mice **(Fig 4A)**. Upon supplementation with oligofructose, the gut microbiota of both lean/control and obese/T2D mice shifted dramatically with the control fiber cellulose have minimal effects **(Fig 4B)**. Discriminant analysis suggested several taxa were altered, including *B. pseudolongum, A. muciniphila*, those from the S24-7 (also known as Muribaculaceae) family, and *Allobaculum* genus **(Fig 4C and 4D)**. Here, we focused on *B. pseudolongum* as an example of the distinct shifts induced by oligofructose because: 1) *A. muciniphila* was not significantly altered in obese/T2D given oligofructose and 2) taxa from the S24-7 family and *Allobaculum* genus are currently uncharacterized. Both S24-7 and *Allobaculum* have been associated with fermentable fiber supplementation and improved glucose tolerance (60, 61). However, our understanding of *A. muciniphila, Allobaculum*, and S24-7 is lacking compared to *Bifidobacterium* spp; thus, our ongoing efforts include isolating and exploring the beneficial effects of these microbes. Overall, oligofructose promoted significant shifts in the gut microbiome of obese/T2D mice when considering both richness and abundance with noteworthy changes in *B. pseudolongum* abundance **(Fig 4D)**.

*Bifidobacterium* spp are well-known producers of the SCFA, immunomodulatory metabolites that are products of fiber fermentation. Similar to previous research that suggests oligofructose induces bifidogenic effects, supplementation with oligofructose led to an expansion in *B. pseudolongum* (**Fig 4D**) (51, 52). Unexpectedly, we did not observe elevated SCFAs in mice given oligofructose as compared to cellulose **(Fig S3)**. However, unique to our study, we observed increased production of the natural polyamines, spermine and spermidine, in plasma and cecum of obese/T2D+OF mice **(Fig 5 and 6)**. Although oligofructose is known to increase SCFAs, quantification is inconsistent across rodent models as duration of supplementation (14 days-6 weeks) and method of isolation varies (feces vs cecal material, fasting vs non-fasting) (35, 51, 62). Short-term treatment with oligofructose (total of 4 weeks) did not affect SCFAs in obese/T2D mice in this study and in an osteoarthritis model from our group (63). Consistent with our metabolomics analysis however is prior documentation of increased polyamine production in rats fed oligofructose for 4 weeks (64). Polyamines are bioactive polycations derived from amino acid metabolism and are essential for cellular proliferation and differentiation. Increasing evidence suggests that polyamines regulate T cell progression, promote macrophage polarization, and reduce production of pro-inflammatory cytokines like TNF-α, which is consistent with our data **(Fig 4)** (53, 65). Polyamines are also involved in osteogenic differentiation, with studies demonstrating exogenous polyamines disrupting osteoclast differentiation leading to attenuated bone loss (66). Therefore, polyamines may be tested to prevent bone destruction associated with chronic infections in obesity/T2D, where mice at 21 days post-infection show significantly more bone loss than lean/control mice (38). We suggest that the changes in the gut microbiota, including increased *B. pseudolongum* abundance, likely leads to increased systemic polyamine production for several reasons: 1) experiments with various *Bifidobacteria* species demonstrated their capability to produce spermidine *in vitro*, 2) shotgun metagenomic sequencing of gut microbiota from obese mice fed resistant starch indicated polyamine synthesis was associated with increases in *B. pseudolongum*, and 3) mice treated with arginine and *Bifidobacterium animalis* led to increased levels of circulating and colonic polyamines, which corresponded with decreased levels of TNF-α and IL-6 (67–69). The evidence from our work demonstrated that polyamines were significantly upregulated in the gut, where microbes are abundant **(Fig 6)**, and the beneficial effects of polyamines on osteomyelitis in obese/T2D mice was observed when administered directly in drinking water **(Fig 7)**. We therefore, conclude that polyamines are immunoregulatory within the context of osteomyelitis rather than directly inhibiting *S. aureus* growth based on our in vitro growth curve results **(Fig S4)**.

In conclusion, we successfully demonstrated the therapeutic potential of the dietary fiber, oligofructose, on osteomyelitis in obese/T2D hosts **(Fig 8)**. Oligofructose reduced bacterial burden, *S. aureus* community colonization in the bone, and ameliorated the hyperinflammatory response in obese/T2D mice, indicating that the fiber reduced infection severity. The observations corresponded with changes in the gut microbiota and overall metabolism that suggested polyamines, and not short-chain fatty acids, were involved in improving infection response. Increased *B. pseudolongum* abundance and the other bacteria affected by oligofructose likely contribute to heightened production of polyamines as their increased abundance was associated with increased acetyl-ornithine, an intermediate of polyamine synthesis produced solely by bacteria. Treating obese/T2D with polyamines in drinking water provided direct evidence for the potential of spermine and spermidine in reducing infection severity. Overall, this study uncovered a novel role for oligofructose and polyamines in the context of bone infections, warranting further investigation into their role in immunoregulation and potential as adjunct therapeutics for obese/T2D hosts against invasive *S. aureus*-mediated osteomyelitis.

## MATERIALS AND METHODS

### Animals

All handling of mice and associated experimental procedures were reviewed and approved by the University Committee on Animal Resources at the University of Rochester Medical Center. Male C57BL/6J mice from The Jackson Laboratory (Bar Harbor, ME) were housed five per micro-isolator cage in a two-way housing room on a 12-hour light/dark schedule. Males were used as they gain weight more consistently than female mice (70). At six weeks of age, mice were provided unrestricted access to either lean (10% kcal fat, OpenSource Diets D12450J) or high-fat (60% kcal fat, OpenSource Diets, D12492) diets for 12 weeks. At week 12, mice were transitioned to supplemented lean and high-fat diets with 10% (w/w) cellulose (Vivapur® 105, JRS Pharma LP, USA) or 10% (w/w) oligofructose (Orafti® P95, BENEO) for two more weeks. All diets were supplied by Research Diets, Inc.; New Brunswick, NJ. For experiments regarding polyamines, mice were continued on either lean or high-fat diets without supplements for the entire duration of the experiment. At week 12, mice were supplemeneed daily with fresh, sterile 3% (total concentration) of spermine (Sigma-Aldrich; St. Louis, MO) and spermidine (Sigma-Aldrich; St. Louis, MO) or PBS in red watter bottles. Bottle weights were measured daily to track drinking habits. All mice were then infected with *S. aureus* at week 14 using a pin model described below and monitored for an additional two weeks following infection. At harvest, mice were sacrificed with CO_2_ and cardiac puncture as a secondary method unless otherwise noted.

### Implant-associated infection model

*S. aureus* USA300 LAC::lux, a kind gift from Tammy Kielian (University of Nebraska Medical Center), was used to infect tibias of mice. Bacteria were added to 10mL tryptic soy broth and cultured overnight at 37°C with shaking at 250 RPM. Stainless steel wire (0.02× 0.5mm) was used to make L-shaped pin implants (4 × 1 mm), which were submerged in overnight culture for 20 minutes prior to implantation (~5 × 10^5^ CFUs). Mice were anesthetized with 60 mg/kg ketamine and 4mg/kg xylazine and received preoperative extended-release buprenorphine. The right tibial area was shaved and disinfected with 70% ethanol and betadine. A ~5 mm incision was used to uncover the tibia, and 30-and 26-gauge needles were used to drill a hole through the bone to fit the pin. Pins were inserted into the drilled holes and the surgical site was closed with 5-0 nylon sutures.

### Glucose tolerance test (GTT)

At week 14 (two weeks after supplementation but prior to infection), mice were fasted for 6 hours during early morning and a glucose tolerance test was performed. Mice were injected intraperitoneally with 300 mg/kg glucose in 0.9% NaCl using U-100 insulin syringes (0.5mL 30G x 5/16) (Beckton Dickinson; Franklin Lakes, NJ). Tail vein blood glucose levels were then monitored before injection and at 15, 30, 90, 120 minutes post-injection using a OneTouch® Verio glucometer and test strips (Lifescan; Milpitas, CA).

### Soft-tissue abscess size and bacterial burden quantification

At harvest, an incision was made at the site of infection to expose soft-tissue abscesses. The shortest and longest diameter of the abscess was measured using a digital caliper (Mitutoyo, Japan), and an average of the two diameters was used as the final measurement for abscess size. For bacterial burden quantification, soft-tissue abscesses and pin implants were homogenized in Lysing Matrix A tubes at 6.0 m/s for 40 seconds with MP FastPrep-24 bead-beating system. Tibias were homogenized for an extra 40 seconds. Homogenized tissue and bone were then serially diluted and plated on tryptic soy agar to determine CFUs.

### μCT and osteolysis analysis

Infected tibias were fixed in 10% neutral buffered formalin at room temperature for 3 days. Samples were then rinsed in PBS and distilled water. Soft tissue and pin implants were removed prior to imaging. Three-dimensional images of infected tibias were acquired by high-resolution micro-computed tomography (vivaCT 40; Scanco Medical AG, Basserdorf, Switzerland). Tibias were scanned with the following parameters: 145 kV energy setting, 300 ms integration time, 10.5 mM voxel size, 10 mM slice increment, and a threshold of 210. Osteolysis was determined by quantification of the void area both from medial and lateral views.

### Histological analysis

Following µCT imaging, fixed tibias were decalcified with 14% EDTA for 7 days. Following decalcification, samples were stored in 70% ethanol until they were processed and embedded in paraffin. Transverse sections of 5µm in size were cut and mounted on glass slides. Images were scanned with VS120 Virtual Slide Microscope (Olympus, Waltham, MA) and visualized using Olympia OlyVIA for abscess counting (minimum of three technical replicates per mouse). Quantification of *S. aureus* communities and TRAP areas were performed using Analysis Protocol Packages in Visiopharm v.2021.07 (Hoersholm, Denmark).

### 16S rRNA sequencing

Samples were processed and sequenced as previously described (63). Fresh fecal samples were collected into sterile microcentrifuge tubes and frozen at −80°C until extraction. Total DNA was extracted from fecal samples using Quick-DNA™ Fecal/Soil Microbe Miniprep (Zymo Research; Irvine, CA) and DNA concentrations were quantified with a nanodrop. V3V4 regions of 16S ribosomal DNA was amplified with PlTaq polymerase (Thermo Fisher Scientific;) using dual indexed primers (319F: 5′ ACTCCTACGGGAGGCAGCAG 3′; 806R: 3′ ACTCCTACGGGAGGCAGCAG 5′). Amplicons were then normalized and pooled using SequalPrep™ Normalization plates (Thermo Fisher Scientific; Waltham, MA). Pooled library was paired-end sequenced on an Illumina MiSeq (Illumina; San Diego, CA) the University of Rochester Genomics Research Center. Each sequencing run included: (1) positive controls consisting of a 1:5 mixture of *Staphylococcus aureus, Lactococcus lactis, Porphyromonas gingivalis, Streptococcus mutans*, and *Escherichia coli*; (2) negative controls consisting of sterile saline; and (3) extraction controls of saline. The background microbiota was monitored at multiple stages of sample collection and processing. All sterile saline, buffers, reagents, and plasticware used for sample collection, extraction and amplification of nucleic acid were UV irradiated to eliminate possible DNA background contamination. Elimination of potential background from the irradiated buffers, reagents, plasticware and swabs was confirmed by 16S rRNA amplification. Data from these background negative control samples is deposited in SRA along with positive controls (BioProject ID: PRJNA786881).

### 16S analysis

Raw reads from the Illumina MiSeq basecalls were demultiplexed using bcl2fastq version 2.19.1 requiring exact barcode matches (71). QIIME 2021.2 was used for subsequent processing (72). 16S primers were removed allowing 20% mismatches and requiring at least 18 bases. Cleaning, joining, and denoising were performed using DADA2 for both runs individually prior to merging. Batch A forward reads were truncated to 265 bps and reverse reads to 244 bps, error profiles were learned with a sample of one million reads, and a maximum expected error of two was allowed. Batch B forward reads were truncated to 246 bps and reverse reads to 216 bps with the same error criteria as Batch A. Batches were merged prior to taxonomic classification with a pretrained naïve Bayesian classifier trained with full-length 16S sequences (73, 74). Diversity analysis was done in R version 4.0.3 using phyloseq_1.32.0 (75). Taxa that could not be classified at least at the phylum level were discarded. Taxa observed in only one sample were also discarded. Samples were rarefied at a depth of 5554 reads, which omitted a total of three samples. Faith’s PD and the Shannon index were used to measure alpha diversity, and Kruskal-Wallis to test for differences. Bray-Curtis dissimilarity index was used to measure beta diversity (76).

### Quantitative PCR

qPCR was used to determine absolute copy numbers of the GroEL gene using primers specific for *Bifidobacterium pseudolongum* (B_plon-std-F 5’ CTCAAGAACGTTGTGGC, B_plon-std-R 5’ CGGTCTTCATCACGAG) (77). Briefly, 2 μL of fecal DNA from each mouse was used as template in 25 μL total reactions of SYBR Green Real Time PCR Master Mix as instructed by the manufacturer (Qiagen; Hilden, Germany). Reactions were carried using CFX Connect Real Time PCR Detection System (BioRad; Hercules, CA) as follows: 95°C 3 min; 40 cycles of 95°C 10 sec, 66°C 30 sec; melt curve 65°C-95°C. A standard curve was made from *B. pseudolongum* ATCC 25526 genomic DNA diluted 10-fold. A trendline was fitted to linear curve and used to calculate absolute copy numbers in fecal samples.

### Multiplexed cytokine array

Blood samples were collected at sacrifice via cardiac puncture and stored in K_2_EDTA anti-coagulant tubes (Beckton Dickinson; Franklin Lakes, NJ). To isolate plasma, blood was centrifuged at 2000 x g for 10 minutes at 4°C. Plasma was transferred to clean microcentrifuge tubes and stored at −80°C until analysis. Plasma samples were analyzed by Eve Technologies Corporation; Calgary, AB, Canada, using a multiplex assay for levels of 31 different cytokines and inflammatory mediators. Cytokines/chemokines with more than one undetectable sample were omitted in heatmap.

### Targeted metabolomics

A separate cohort of mice was dedicated to targeted metabolomic profiling. Mice were fed on the same feeding scheme as before, but were harvested two weeks after supplementation without infection to prevent confounding effects of infection on metabolism. Mice were fasted for six hours prior to intraperitoneal injection with 100mg/kg Euthasol (phenytoin/pentobarbital) and cervical dislocation. Plasma was isolated as previously described from blood collected via cardiac punctures. Cecal material was isolated using sterile forceps and placed directly into clean, pre-weighed microcentrifuge tubes. Both plasma and cecal samples were snap-frozen in liquid nitrogen and stored at −80°C prior to metabolomic analysis. The Metabolomics Innovation Centre (Alberta, CA) was contracted to identify and quantify up to 150 metabolites commonly found in samples.

A targeted quantitative metabolomics approach was used to analyze the samples using a combination of direct injection mass spectrometry with a reverse-phase LC-MS/MS custom assay. This custom assay, in combination with an ABSciex 4000 QTrap (Applied Biosystems/MDS Sciex) mass spectrometer, can be used for the targeted identification and quantification of upto 150 different endogenous metabolites including amino acids, acylcarnitines, biogenic amines & derivatives, uremic toxins, glycerophospholipids, sphingolipids and sugars (78, 79). The method combines the derivatization and extraction of analytes, and the selective mass-spectrometric detection using multiple reaction monitoring (MRM) pairs. Isotope-labeled internal standards and other internal standards are used for metabolite quantification. The custom assay contains a 96 deep-well plate with a filter plate attached with sealing tape, and reagents and solvents used to prepare the plate assay. First 14 wells were used for one blank, three zero samples, seven standards and three quality control samples. For all metabolites except organic acid, samples were thawed on ice and were vortexed and centrifuged at 13,000x g. 10 µL of each sample was loaded onto the center of the filter on the upper 96-well plate and dried in a stream of nitrogen. Subsequently, phenyl-isothiocyanate was added for derivatization. After incubation, the filter spots were dried again using an evaporator. Extraction of the metabolites was then achieved by adding 300 µL of extraction solvent. The extracts were obtained by centrifugation into the lower 96-deep well plate, followed by a dilution step with MS running solvent. For organic acid analysis, 150 µL of ice-cold methanol and 10 µL of isotope-labeled internal standard mixture was added to 50 µL of sample for overnight protein precipitation. Then it was centrifuged at 13000x g for 20 min. 50 µL of supernatant was loaded into the center of wells of a 96-deep well plate, followed by the addition of 3-nitrophenylhydrazine (NPH) reagent. After incubation for 2h, BHT stabilizer and water were added before LC-MS injection. Mass spectrometric analysis was performed on an ABSciex 4000 Qtrap® tandem mass spectrometry instrument (Applied Biosystems/MDS Analytical Technologies, Foster City, CA) equipped with an Agilent 1260 series UHPLC system (Agilent Technologies, Palo Alto, CA). The samples were delivered to the mass spectrometer by a LC method followed by a direct injection (DI) method. Data analysis was done using Analyst 1.6.2.

## List of Abbreviations

Obesity/T2D: obesity-related type 2 diabetes
OF: oligofructose
CL: cellulose
qPCR: quantitative polymerase chain reaction
LC-MS: liquid chromatography-mass spectrometry
DI: direct injection
CFU: colony forming unit
PBS: phosphate buffered saline
SCFA: short-chain fatty acids
PA: polyamine
GTT: glucose tolerance test

## Availability of Data and Materials

Illumina 16S rRNA V3V4 amplicons were deposited in Sequence Read Archive under BioProject PRJNA786881, including positive and negative controls on each plate.

## Author Contributions

TIB, RAM, and SRG designed the study. TIB and ALG carried out all experimental studies and generated data. TIB analyzed all data. TIB, RAM, and SRG drafted the manuscript. All authors reviewed the final manuscript.

## Acknowledgements

We thank the Center for Musculoskeletal Research for providing procedure space, as well as Jeffrey Fox for his technical contributions with histological analysis and Lindsay Schnur for her contributions with micro-CT. We also thank the Genomics Research Center for performing16S rRNA sequencing and Cal Palumbo for his guidance with taxonomic analysis.

## SUPPLEMENTAL FIGURE LEGENDS

**Figure S1. Diet-induced obese/T2D mice exhibit hyperglycemia compared to lean/control mice.** After two weeks of supplementation with cellulose or oligofructose (week 14), mice were fasted six hours prior to glucose tolerance testing. Blood glucose levels were monitored using OneTouch Verio glucometer on blood from tail veins prior to injection with a bolus of glucose (300mg/kg). Blood glucose levels were then measured at time points = 15, 30, 60, and 120 minutes. Bar graphs represent mean ± SD. n=5. Significance was identified using one-way ANOVA and Tukey’s post-hoc multiple comparisons test. *\*\*P*<0.01, *\*\*\*P*<0.001, *\*\*\*\*P*<0.001.

**Figure S2. Top 12 genera of gut microbiota from fecal samples.** A) Week 12, prior to supplementation. B) Week 14, two weeks post-supplementation with fiber but prior to infection. C) Week 16, two weeks after infection while supplemented with fiber. NS=No supplement; CL=Cellulose (control fiber); OF=Oligofructose.

**Figure S3. Oligofructose does not alter SCFA concentrations in plasma or cecum of obese/T2D mice.** Short-chain fatty acids (SCFAs) were quantified by targeted metabolomics on samples isolated after two weeks of supplementation without infection as previously described. Bar graphs represent mean ± SD. n=5. Significance was identified using one-way ANOVA and Tukey’s post-hoc multiple comparisons test. *\*\*P*<0.01, *\*\*\*P*<0.001, *\*\*\*\*P*<0.001.

**Figure S4. *S. aureus* growth in the presence of polyamines.** A) *S. aureus* USA300 was incubated with various concentrations of spermine and spermidine in a 96-well plate and absorbance at 600 nm was monitored over 24 hours. SP (spermine), SPD (spermidine), TSB (tryptic soy broth) B) Area under curve was used to quantified overall growth across groups. Bar graphs represent mean ± SD. n=3. Significance was identified using one-way ANOVA and Tukey’s post-hoc multiple comparisons test.

